# Optimisation and comparison of orthogonal methods for separation and characterisation of extracellular vesicles to investigate how representative infant milk formula is of milk

**DOI:** 10.1101/2020.07.27.221622

**Authors:** Anindya Mukhopadhya, Jessie Santoro, Barry Moran, Zivile Useckaite, Lorraine O’Driscoll

## Abstract

This study aimed to separate and characterise extracellular vesicles (EV) from infant milk formula (IMF) and skim milk (SM), to determine how representative the EV content of IMF is to SM. Contaminant casein micelles, due to their abundance and overlapping size, were removed followed by either differential ultracentrifugation (DUC) or gradient ultracentrifugation (GUC). Characterisation included BCA, SDS-PAGE, nanoparticle tracking analysis (NTA), transmission electron microscopy (TEM), immunoblotting, and imaging flow cytometry (IFCM). NTA reported significantly reduced concentrations of EVs/particles in IMF versus SM; TEM showed intact SM-derived EVs to sparse and disrupted EV-like structures in IMF. Compared to IMF, noticeably stronger bands for EV biomarkers were observed by immunoblotting in SM, indicating compromised EV proteins in IMF; also supported by IFCM. Altogether, we established that EVs are substantially compromised during IMF processing. Furthermore, an optimised method combining acid pre-treatment and GUC for EV separation from milk products has been established.

## 1. Introduction

Bovine milk is considered as a major source of nutrition for humans and consumed, across the globe, by all age groups. The proposed biological function of extracellular vesicles (EVs) in milk, including their role as cell growth and immune system modulator, is gathering increasing interest (Benmoussa & Provost, 2019). Infant milk formula (IMF), typically made from bovine skim milk (SM), is frequently used to fulfil the nutritional requirement of infants and toddlers. However, although the industrial processing stages of IMF manufacture from SM involves extensive thermal treatments steps - including sterilisation, evaporation and spray-drying that can involve temperatures of up to 280°C (Burke et al., 2018; Gathercole et al., 2017)- the influences of these harsh conditions on EVs within IMF has not been addressed. This may be, at least in part, due to the fact that milk and IMF are very complex fluids and a suitable method that is compliant with MISEV2018 for separating and characterising EVs from both has not yet been reported. However, as the EU Commission Directive (2006/141/EC) specifies that IMF composition must be of sufficient biological quality, IMF and dairy ingredient industries needs to know how representative IMF is of SM. They need to know whether EVs are present and maintained as in SM or, alternatively, if efforts would be needed to retain/supplement EVs. Simply put, industry -and consumers-need to know what’s in the IMF product.

Substantial efforts have been invested in analysing EVs from bovine milk (Arntz et al., 2015; Hata et al., 2010; Munagala et al., 2016; Reinhardt et al., 2013; Samuel et al., 2017; Somiya et al., 2018; Vaswani et al., 2017; Wolf et al., 2015; Yamada et al., 2012) and human milk (Admyre et al., 2007; Lasser et al., 2011; M. J. C. van Herwijnen et al., 2018; Zonneveld et al., 2014). The removal of any present debris and milk protein casein before proceeding to EV separation was considered; studies on bovine milk have applied either differential centrifugation or treatment with EDTA or acids, whereas those on human milk performed mostly differential centrifugation and filtration. However, till date, a standardised pre-analytical protocol for casein removal seems to be lacking, considering the extremely high concentrations 10^14^–10^16^/mL of casein in milk (Fox, 2001; O’Regan et al., 2009) that, because of their size (approx. 150 nm), fall well within the size range of EVs. From food science point of view, casein and casein micelles are important due to their various applications and caseins are separated from milk by either adjusting milk pH to ∼4.6 (isoelectric precipitation point) using acids or by ultracentrifugation (Jensen et al., 2012).

Based on a survey in 2016, differential ultracentrifugation (DUC) is the most commonly used EV separation technique and alternative techniques such as density gradient ultracentrifugation (GUC), size exclusion chromatography, and precipitation techniques being used to a varying degree (Gardiner et al., 2016). Since the first study was published by Admyre et al., (2007) on the presence of EVs in breast milk, separated by DUC, there was an increase in studies using DUC step as the primary and/or additional method for isolation of EVs was observed (Izumi et al., 2012; Kusuma et al., 2016; Manca et al., 2018; Reinhardt et al., 2012; Q. Sun et al., 2013). Interestingly, as reviewed by Witwer et al., (2013), pure EV samples are not separated by DUC methods as DUC does not take into consideration the density of the EV cargo, whereas, GUC, especially the ‘bottom-up’ approach, efficiently separates the contaminant proteins from EVs. Since then, milk EV studies have started considering GUC as the main method of EV separation with or without additional purification methods (Benmoussa et al., 2016; J. Sun et al., 2015; M. J. van Herwijnen et al., 2016; Yamauchi et al., 2018a).

Hence, this current study compared EVs from SM and IMF, separated by GUC and DUC after investigating options of pre-treatment (acetic acid or isoelectric precipitation) to remove casein micelles that would otherwise confuse EV analysis. Resulting isolates were then compared based on their particle content, protein content and EV surface markers. We hypothesise that industrial processing leads to disruption of EV present in the milk. We also hypothesise that using only ultracentrifugation technique is not enough to remove the casein micelles from milk and hence a pre-treatment is essential to separate pure EVs from milk. A graphical representation of the different milk treatments, EV separation methods and characterisation methods used is summarised in Figure 1.

**Figure 1.**
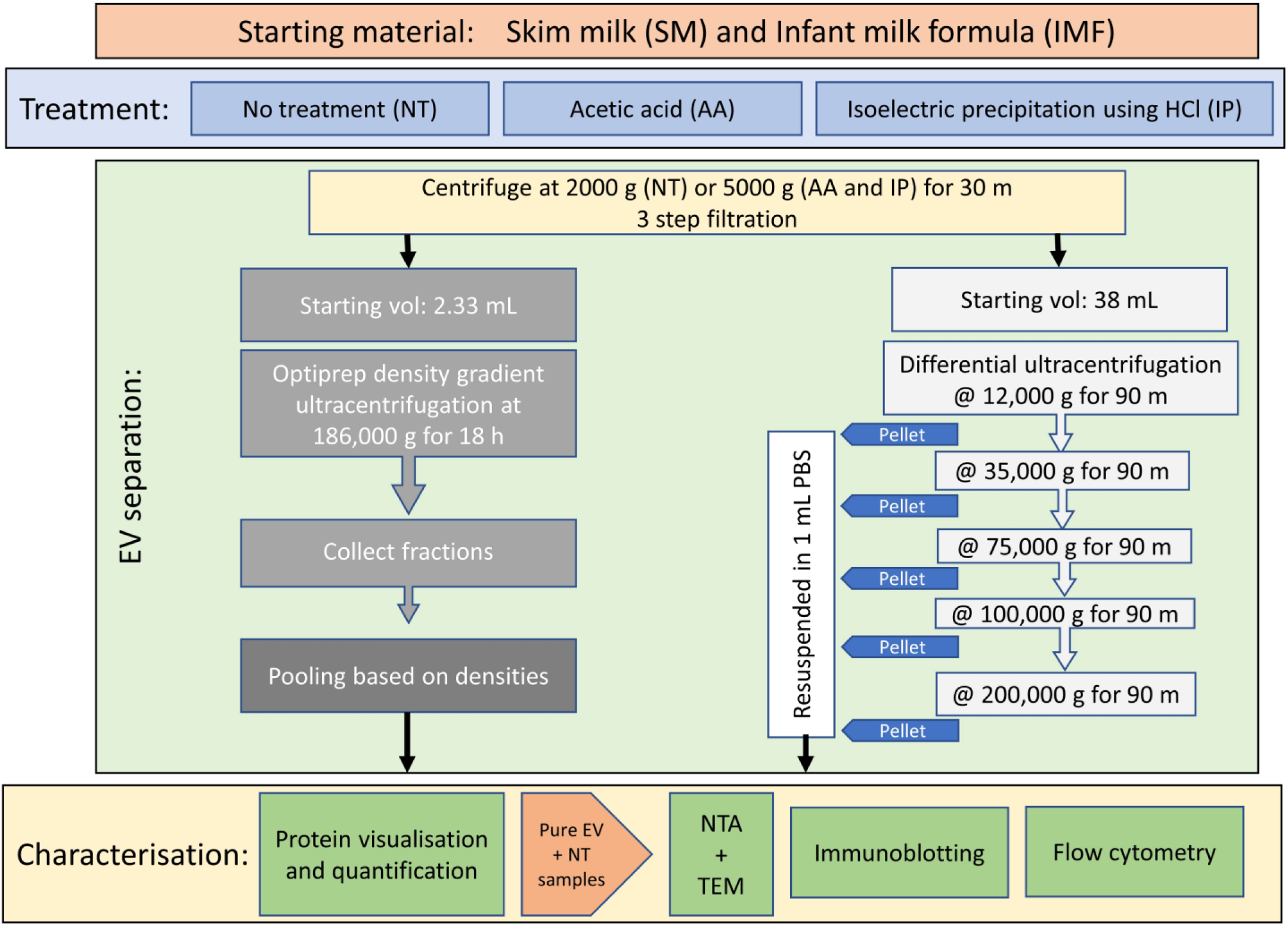
Overview of methodology used. A flow-chart of the methodology used in EV separation from skim milk (SM) and infant milk formula (IMF). The flow-chart is divided into sections including the starting materials used, treatments performed, ultracentrifugation techniques used and characterisation of the samples.

## 2. Materials and Methods

All relevant information was submitted to EV-TRACK knowledgebase (Van Deun et al., 2017) The EV-TRACK ID: EV190096 was assigned and the score achieved was 100%.

### 2.1 Milk samples

Fresh batches (n=3 each) of bovine skim milk and infant milk formula (IMF) prepared as recommended by the manufacturer were purchased from local Irish supermarket and used throughout. Each batch of SM and IMF were divided into 3 equal aliquots. One was untreated (NT), and the others were to be treated with either acetic acid (AA) or HCl for isoelectric precipitation (IP) in an effort to remove casein micelles. All subsequent analyses detailed in the study were performed a minimum of 3 times.

### 2.2 Filtration and treatment for casein micelles removal

For the untreated samples, 50 mL of SM or IMF were centrifuged at 2,000 g for 20 min at 4°C. The supernatant was carefully collected and the pellet containing any remaining fat globules or debris was discarded. This supernatant then underwent a 3-step filtration i.e. it was passed through Whatman™ Grade 1 filter paper (diameter 150 mm) (GE Healthcare Life Sciences, Ireland), followed by 0.45-μm membrane filtration and then 0.22-μm membrane filtration (ThermoFisher Scientific Ltd). The filtrate obtained from untreated SM was designated as SM_NT and the filtrate obtained from untreated IMF was designated as IMF_NT.

The AA-treatment of SM and IMF was performed using a modification of a method previously reported for defatted bovine milk (Somiya et al., 2018). However, here additional filtration and ultracentrifugation steps were included in an effort to remove all casein protein aggregates and casein micelles, which was not completely achieved as reported (Somiya et al., 2018). In brief, SM or IMF was mixed with acetic acid (Sigma-Aldrich, Ireland) [milk/acetic acid = 100% (v/v)] for 10 min at RT and centrifuged at 5,000 g for 30 min at 4°C. Here, instead of performing a 0.22-μm filtration and ultracentrifugation of the resulting whey at 210,000 g, the 3-step filtration (detailed above) was performed prior to subsequent differential ultracentrifugation. The filtrate from SM was designated as SM_AA and that from IMF was designated as IMF_AA.

Isoelectric precipitation (IP) was performed using a modification of a method previously reported by Yamauchi et al. (2018b) for raw bovine milk. In brief, here the pH of SM or IMF were adjusted to pH4.6 using 6 N HCl (Sigma-Aldrich, Ireland) to remove caseins. The pH adjusted SM or IMF was placed on ice for 10 min followed by centrifugation at 5,000 g for 30 min at 4°C. Resulting whey was subject to the 3-step filtration procedure. The filtrate from SM was designated as SM_IP and that from IMF was designated as IMF_IP. While the protocol used by Yamauchi et al. (2018b) then proceeded to a sucrose gradient we, instead, use OptiPrep™ gradient and differential ultracentrifugation techniques, detailed in the following sections.

Of note, neither Somiya et al. (2018) nor Yamauchi et al. (2018b) analysed IMF.

### 2.3 Separation of EVs by ultracentrifugation (UC) using either Gradient UC or Differential UC

Gradient ultracentrifugation (GUC) was performed using iodixanol solution OptiPrep™ (60% (w/v)) (Sigma Aldrich, Cat. #D1556) following the bottom-up technique (Kowal et al., 2016). Briefly, a 40% (w/v) bottom layer was made by diluting 60% Optiprep solution with each of the 6 sample types (4.66 mL Optiprep + 2.33 mL sample = 7 mL). Further layering was done by a discontinuous gradient of 30% (w/v), 20% (w/v), 10% (w/v) and 5% (w/v) solutions of iodixanol, made by diluting a stock solution of OptiPrep (60% (w/v)) with sterile PBS (Sigma Aldrich, Ireland). The gradient was formed by adding 3 mL of the iodixanol solutions to a 17 mL polypropylene tube (Beckman Coulter). Ultracentrifugation was performed using a SW Type 32.1 Ti rotor in Optima XPN-100 Ultracentrifuge (Beckman Coulter, Brea, CA, USA) at 186,000g for 18 h at 4 °C. Post 18 h centrifugation, the supernatant was carefully pipetted out in 1 mL fractions. Their refractive indexes were recorded, and densities were calculated for all fractions. Based on previous studies (Colombo et al., 2014; Kowal et al., 2016), in this study, fractions with densities of 1.05-1.19 g/mL were pooled for further analyses. Of note, as we aimed to isolate all EVs present in SM and IMF and considering that Kowal et al. (2016) observed positive signals for EV-specific proteins in fractions with densities of 1.08 g/mL, we chose 1.05–1.19 g/mL fractions. Post-pooling, the sample was washed in sterile PBS at 110,000 g for 90 mins at 4 °C before resuspending the pellet in 1 mL of sterile PBS.

Differential ultracentrifugation (DUC) separation of EVs from the untreated or treated SM and IMF samples was performed based on protocol published by Benmoussa et al. (2017) with addition, in this study, of a final step of ultracentrifugation at 200,000g. A similar separation of EVs from urine (Gonzales et al., 2010; Musante et al., 2013) and milk (Somiya et al., 2018) at 200,000g have been reported previously. In brief, samples were filled into 38 mL open-top polypropylene tubes and subjected to DUC using a SW Type 32 Ti swinging bucket rotor (Beckman Coulter) in an Optima XPN-100 Ultracentrifuge (Beckman Coulter, Brea, CA, USA). Consecutive spins were at 12,000 g (12K), 35,000 g (35K), 75,000 g (75K), 100,000 g (100K) and 200,000 g (200K) for 90 min each at 4 °C. After each centrifugation step, the resulting pellets (12K, 35K, 75K, 100K or 200K) were collected, resuspended in 1 mL filtered PBS and stored in −80 °C for further analysis, while the supernatant was poured in a fresh tube and centrifuged at the next speed.

### 2.4 SDS-PAGE for protein visualisation

Effects on milk proteins of pre-treatment with AA or IP compared to NT, prior to EV separation by ultracentrifugation, was visualised on a 10% polyacrylamide gel (Bio-Rad Laboratories). For this, 15 µg of protein samples were loaded on a 10% polyacrylamide gel, run at 120 volts, stained with coomassie blue (Sigma Aldrich, Ireland) and visualised using a Chemidoc XRS system (Bio-Rad).

### 2.5 Protein quantification

A bicinchoninic acid (BCA) reagent kit (Fisher Scientific, Cat. #10249133), with a bovine serum albumin (BSA) standard (0–800 μg/mL) (Sigma Aldrich, Ireland) curve, was used to determine total protein in samples.

### 2.6 Nanoparticle Tracking Analysis

Particle concentration and sizes were estimated using NanoSight NS300 (Malvern Instruments Ltd, Malvern, UK). EV samples were automatically injected into the NTA system under constant flow conditions (flow rate = 50) and videos of the particles in motion were recorded and analysed using NTA 3.1.54 software.

### 2.7 Transmission electron microscopy

Samples were prepared for transmission electron microscopy (TEM) analysis following previously published protocol (Visnovitz et al., 2019). Specifically, 5 μL of sample was suspended in 5 μL of 0.22μm-filtered PBS and placed onto carbon coated grids (Ted-Pella B 300M, Mason Technology LTD, Cat. #: 01813-F) for 10 minutes at RT before fixing with 4% glutaraldehyde (Sigma Aldrich) and contrasting with 2% phosphotungstic acid (Sigma Aldrich). The grids were examined at 100 kV using a JEOL JEM-2100 TEM (JOEL, Peabody, USA), as we previously described (O’Brien et al., 2013).

### 2.8 Immunoblotting

Immunoblotting was performed as we previously described (O’Brien et al., 2013), except that 35 μg of protein was used here. For this, EVs were lysed using SDS lysis buffer (250 nM Tris-HCL, pH 7.4; 2.5% SDS), lysates (35 μg) were resolved on 10% SDS gels (Bio-Rad) and the protein was transferred onto PVDF membranes (Bio-Rad, Cat. #1620177). Blots were blocked with 5% (w/v) BSA in PBS containing 0.1% Tween-20 and incubated overnight at 4°C with primary antibodies to TSG101 (1:1000; Abcam, Cambridge, UK, Cat. # ab30871), CD63 (1:500; Abcam, Cat. #: ab68418) or Actinin 4 (1/1000, Abcam, Cat. #: ab41525). Secondary antibodies used were anti-mouse (1/1000 in 5% BSA/PBS-T, Cell Signalling, Cat. #: 7076) or anti-rabbit (1/1000 in 5% BSA/PBS-T, Cell Signalling, Cat. #: 7-74). SuperSignal West Femto Chemiluminescent Substrate Kit (Fisher Scientific, Cat. # 11859290) was used for detection, imaging was performed using an automated Chemidoc exposure system (Bio-Rad Laboratories) and densitometric analysis was performed using ImageJ tool.

### 2.9 Imaging flow cytometry

The imaging flow cytometry (IFCM) analysis of EVs separated from SM and IMF was performed following a protocol previously published in JEV (Ricklefs et al., 2019). Briefly, EVs surface antigens were exposed to antibodies diluted in 0.22 μm-filtered PBS with 2% EV-depleted-FBS supplemented with protease inhibitor and phosphatase inhibitor (IFCM buffer). The specific antibodies used were anti-CD63 conjugated with FITC (CD63-FITC; 1:150, Cat. #: 353006, Biolegend); CD9-PE (1:1500, Cat. #: 312106, Biolegend); CD81-PE-Cy7 (1:150, Cat. #: 349512 Biolegend); HLA-DR-BV421 (1:600, Cat. #: 307636, Biolegend) and ADAM10-APC (1:150, Cat. #: 352706, Biolegend). The selection of the antibodies was made based on the presence of surface markers on the EVs. Anti-CD9, anti-CD63 and anti-CD81 antibodies were selected as EV specific tetraspanins present on their surface, along with HLADR, an MHC II immunoregulatory molecule present on EV surface. Additionally, ADAM10 was also selected as in a previous study, Kowal et al. (2016) reported the presence of ADAM10 in EVs. The EVs were incubated with the antibodies for 45 min at RT in the dark, and washed using a 300 kDa filter (Nanosep, Cat. #: 516-8531), resuspended in 50 μl IFCM buffer and acquired within 2 h on the ImageStream X Mk II imaging flow cytometer (Amnis/ Luminex, Seattle, USA) at 60x magnification and low flow rate, as previously described (Ricklefs et al., 2019). EV-free IFCM buffer, unstained EVs, single-stained controls and fluorescence minus one (FMO) controls were run in parallel. Fluorescence was within detection linear range in the following channels: FITC was measured in channel 2 (B/YG_480–560 nm), PE in channel 3 (B/YG_560–595 nm filter), PE-Cy7 in channel 6 (B/YG_745-780 nm filter), BV421 in channel 7 (R/V_435-480 nm filter), APC in channel 11 (R/V_642–745 nm, filter), Brightfield in channel 1 and 9 (B/YG_435-480 and R/V_560-595 nm filter respectively) and Side Scatter Channel (SSC) in channel 12 (R/V_745-780 nm filter). Data analysis was performed using IDEAS software v6.2 (Amnis/ Luminex, Seattle, USA). EVs were gated as SSC-low vs fluorescence, then as non-detectable brightfield (fluorescence vs Raw Max Pixel Brightfield channel), gated EVs were confirmed in the IDEAS Image Gallery.

### 2.10 Statistical analysis

All data presented are in form of the means of ≥3 experiments ± SEM. Unpaired two-tailed Student’s t-test was used to compare two groups and for more than two groups one-way or two-way ANOVA analysis were performed using Graph Pad Prism 8 software. A value of P < 0.05 was considered statistically significant.

## 3. Results

### 3.1 Analysis of milk protein removal by acid treatment and ultracentrifugation

Total protein separation by polyacrylamide gel showed that the major groups of milk proteins, including caseins and whey proteins, were present at high levels in the SM_NT samples post-GUC and to a lesser extent in IMF_NT samples (Fig. 2a i), while treatment with AA- or IP-efficiently removed caseins. Post-DUC, the SM_NT and IMF_NT gave stronger signals for caseins and whey proteins (Fig. 2a ii), while AA- and IP-treatment efficiently remove the casein and whey proteins bands.

**Figure 2.**
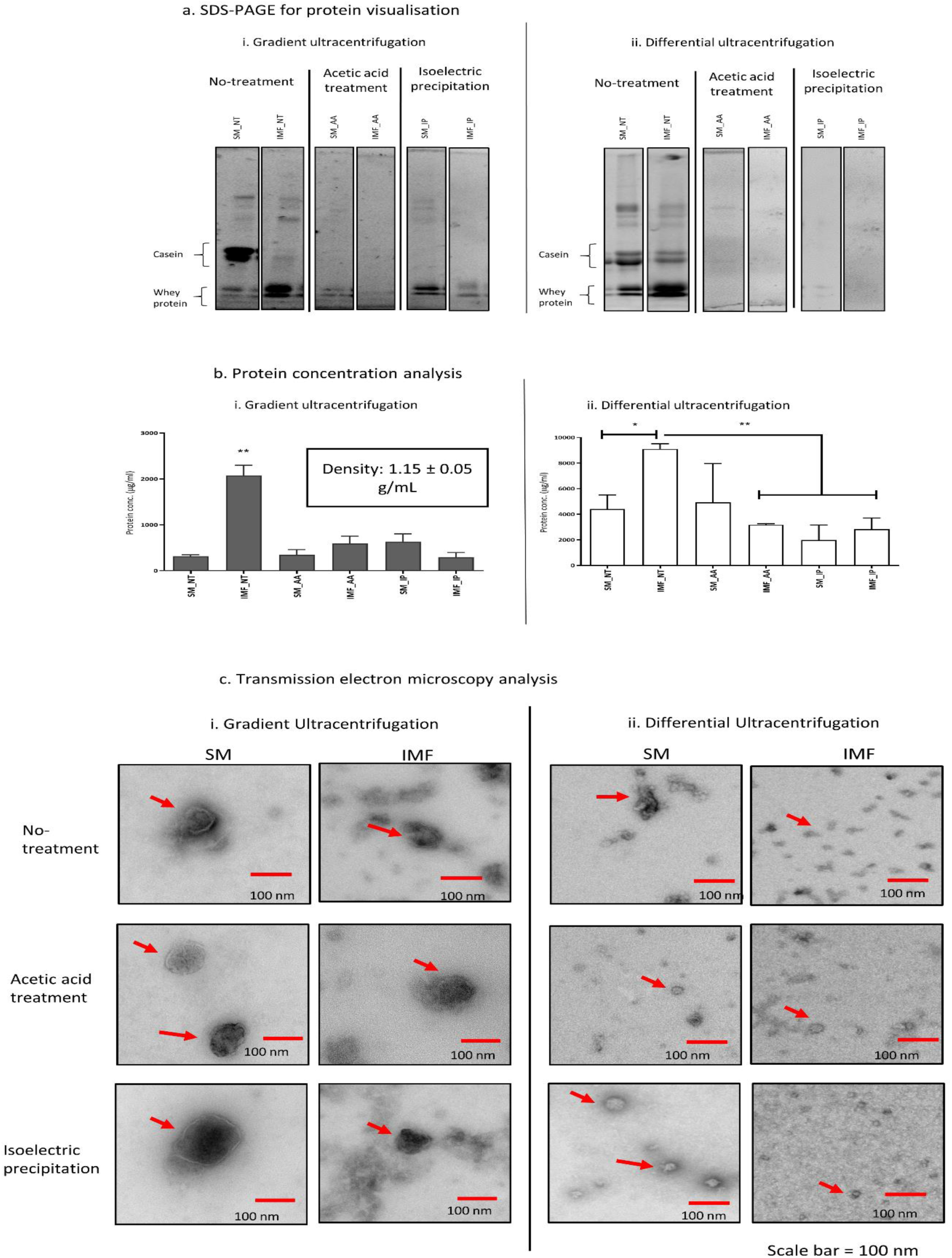
Protein and particle characterisation of skim milk and infant milk formula Evs. a) Milk proteins were visualised on 10% acrylamide gels. The removal of milk proteins was performed by acetic acid (AA) and isoelectric precipitation (IP) and the whey was used for either gradient ultracentrifugation (GUC) or differential ultracentrifugation (DUC). Untreated milk (NT) was used to compare the removal of milk proteins by acid treatment. i) Post GUC, equal amount of protein samples (15 µg) were loaded and electrophoresis performed until separation was achieved for untreated SM and IMF samples (NT), SM and IMF samples collected post AA and SM and IMF samples collected post IP are presented. ii) post-DUC, equal amount of protein samples (15 µg) were loaded and electrophoresed until clear separation was obtained, the 200K resuspended pellet of untreated SM and IMF samples, SM and IMF samples collected post AA and SM and IMF samples collected post-IP are presented. b) Isolates from skim milk (SM) and infant milk formula (IMF) post i) gradient ultracentrifugation (GUC) and ii) differential ultracentrifugation (DUC) were analysed for protein concentrations using bicinchoninic acid (BCA). Protein concentration (in µg/mL) of the three repeats were analysed using one-way Anova test and presented as mean bars ± SEM, * indicates a significance value of P < 0.05, ** indicates a significance value of P < 0.01. c) Representative transmission electron microscope (TEM) images from gradient ultracentrifugation (GUC) and differential ultracentrifugation (DUC) samples. i) TEM micrographs of EV from skim milk (SM) and infant milk formula (IMF) samples collected post GUC, ii) TEM micrographs of samples collected post DUC. Red arrows point towards vesicular structures in the sample. Scale bar=100nm

Due to the presence of casein bands, the DUC generated 12K, 35K, 75K and 100K resuspended pellets or the crude EV samples were not used for further extensive analysis, whereas the 200K samples - were further characterised (representative SDS-PAGE blots on all samples are provided in Supplementary information, Fig. S1 a&b). It is also important to note that in the untreated SM and IMF samples, post-GUC or after 200,000 g ultracentrifugation step, casein and whey proteins are still present to varying extents and so will be present during further characterisation steps as these samples serve as control samples for further characterisation steps.

### 3.2 Protein concentration analysis of isolates from untreated or AA- or IP-treated SM and IMF

The total protein concentrations of samples post-GUC and -DUC are presented in Fig. 2b. Considering the pooled fractions (1.05-1.19 g/mL), the total protein content of IMF_NT appeared to be significantly (P<0.01) higher compared to all other treatment samples analysed (Fig. 2b i).

The total protein contents on post-DUC samples are presented in Fig. 2b ii. The IMF_NT samples were associated with significantly higher total protein content compared to SM_NT samples (P<0.05) and IMF_AA, SM_IP and IMF_IP samples (P<0.01).

### 3.3 Milk EV concentration and size characterisation

The concentrations and sizes of EVs separated from treated or untreated SM and IMF, analysed by NTA, are presented in Table 1. The size distribution of the post-GUC particles from the SM samples were 156.13±5.87 nm and from IMF samples were 159.90±12.96 nm; those post-DUC of SM_200K were 159.10±5.27 nm and IMF_200K were 175.73±8.44 nm. No significant differences were observed between these EV sizes. The representative size distribution graphs with mean and modal sizes (as observed with the NTA software) are also provided in Supplementary information Table S1 a&b.

Regarding the concentration of particles in each sample, significant differences were observed between SM and IMF samples. As presented in Table 1a, post-GUC, the concentration of EVs recorded in SM_NT samples (4.35×10^10^±1.40×10^10^ particles/mL of SM) were significantly (P<0.05) higher than IMF_NT samples (6.13 ×10^09^±8.17×10^08^ particles/mL of IMF). The concentration in SM_AA samples were 1.39×10^10^±8.25×10^09^ particles/mL of SM and IMF_AA concentration was 2.82×10^09^±2.32×10^09^ particles/mL of IMF; statistical significance was not achieved (P = 0.26). And finally, the EV concentration of SM_IP samples (2.16×10^10^±2.47×10^09^ particles/mL of SM) were significantly (P < 0.01) higher compared to IMF_IP samples (3.23×10^09^±1.45×10^09^ particles/mL of IMF).

Similar significant differences in concentration of particles in the suspension resulting post-DUC were also observed, as presented in Table 1b. The concentration of SM_NT_200K samples (1.97×10^10^± 8.93×10^09^ particles/mL of SM) were significantly (P<0.05) higher than IMF_NT_200K samples (4.66×10^07^±2.47×10^07^ particles/mL of IMF) Similarly, the concentration of particles in SM_AA_200K samples (2.77×10^09^±1.04×10^09^ particles/mL of SM) was significantly (P<0.05) higher compared to IMF_AA_200K (1.69×10^08^ ±3.28× 10^07^ particles/mL of IMF). Also, the concentration of particles in the SM_IP_200K samples (3.65×10^09^±5.42× 10^08^ particles/mL of SM) was significantly (P<0.05) higher compared to IMF_IP_200K samples (4.66×10^08^±2.02×10^08^ particles/mL of IMF).

Complementary information was obtained by TEM and presented in Fig. 2c. Following GUC separation, the SM EVs (SM_NT, SM_AA and SM_IP) appeared intact with limited background debris. IMF-derived EVs appeared less clear, with particles of less smooth surfaces, and with a background of debris (Fig. 2c). In the post-DUC SM_NT, SM_AA and SM_IP samples, multiple smaller particles were observed (arrows, Fig. 2c) while the IMF samples (IMF_NT, IMF_AA and IMF_IP) had few, if any, EV like structures.

### 3.4 Immunoblotting analysis of EV specific positive and negative markers

In line with MISEV2018 guidelines (Thery et al., 2018), immunoblotting was done on lysed samples for two EV positive markers, TSG101 and CD63, and a negative marker Actinin 4. The GUC generated SM_NT, SM_AA and SM_IP samples were typically positive for both TSG101 and CD63 and no signals obtained for Actinin 4 (Fig. 3a; densitometric analysis of n=3 presented in Fig. 3b). Interestingly, compared to SM samples, TSG101 and CD63 were detected to varying degrees in IMF_NT, IMF_AA and IMF_IP samples; no signals obtained for Actinin 4 (Fig. 3a).

**Figure 3.**
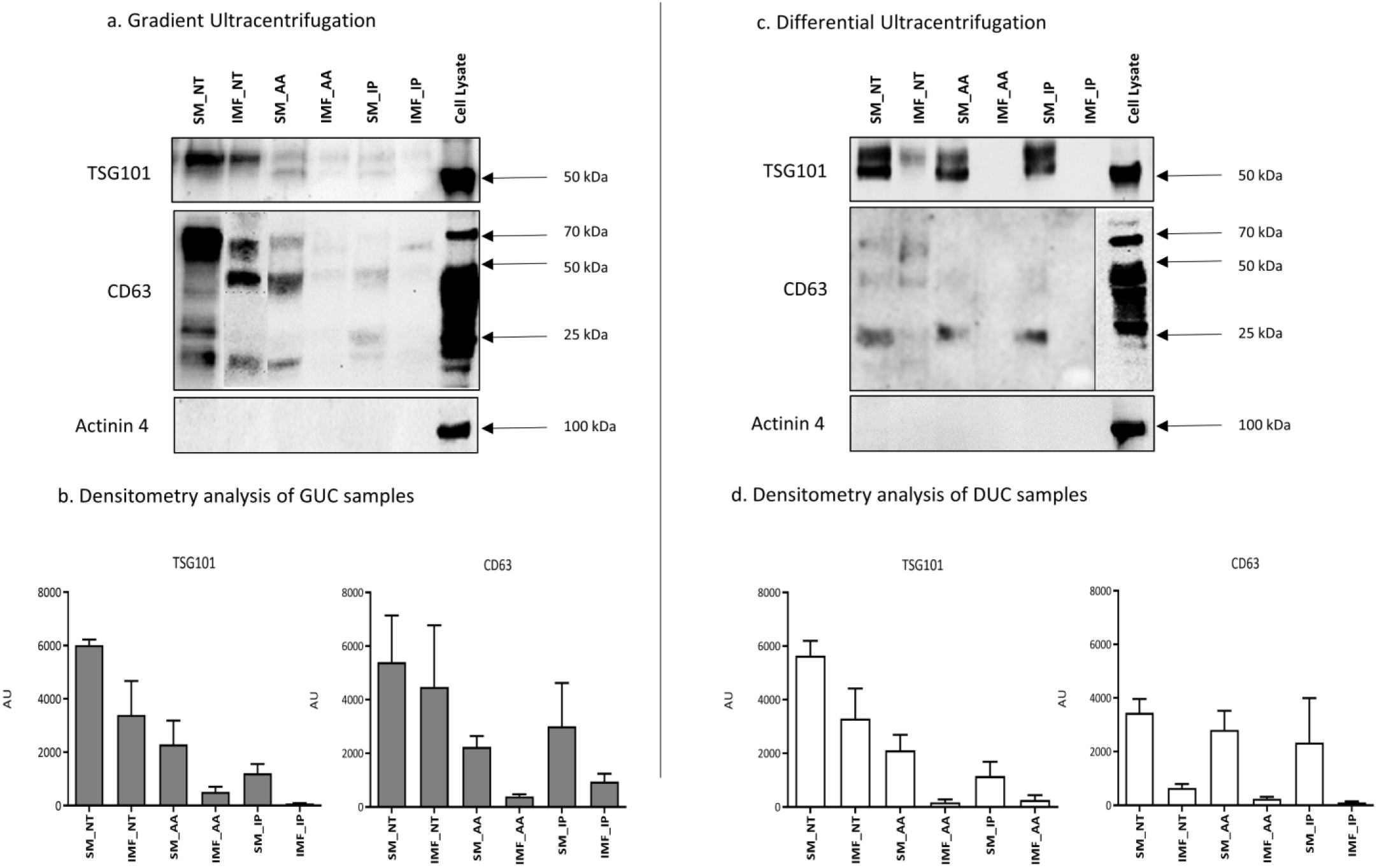
Immunoblotting of EV specific markers. Immunoblotting of skim milk (SM) and infant milk formula (IMF) EVs separated using gradient ultracentrifugation (GUC) and differential ultracentrifugation (DUC) and their densitometric analysis. Equal amount of protein (35 µg) were loaded for all the samples, electrophoresed until optimum separation was achieved and then transferred on to a PVDF membrane before staining with primary (overnight) and secondary (1 h) antibodies before imaging using hi-sensitivity chemiluminescence under Chemidoc XRS system. Representative blots for GUC samples are presented in (a) and DUC samples are presented in (c), overall densitometric analysis of GUC samples are presented in (b) and those of DUC samples in (d). ImageJ software was used to perform densitometric analysis. Each bar represents the mean of the densities of the signals obtained post immunoblotting (mean of n=3 biological repeats ± SEM).

All the EVs/particles collected post-DUC of SM and IMF samples were also lysed and analysed for TSG101, CD63 and Actinin 4 (Fig. 3c and densitometric analysis of n=3 presented in Fig. 3d). No signals for Actinin 4 were observed for both SM and IMF samples generated by DUC. SM_NT, SM_AA and SM_IP samples were positive for both TSG101 and CD63. IMF_NT samples compared to SM_NT had faint TSG101 and CD63 signals. Interestingly, both IMF_AA and IMF_IP samples compared to SM_AA and SM_IP samples respectively, showed no bands for both TSG101 and CD63.

### 3.5 Detection of EVs using Imaging flow cytometry analysis

Detection of EVs based on presence of EV surface markers CD9, CD63, CD81, ADAM10 and HLADR was performed using imaging flow cytometry (IFCM). Based on a recent JEV publication (Ricklefs et al., 2019) we have provided the gating strategy used in Supplementary Fig. S2 and representative EV images from IDEAS Image Gallery in Supplementary Fig. S3.

Post-GUC (Fig. 4a), significant differences in the particles/mL of SM_AA *vs*. IMF_AA (P < 0.01) and SM_IP *vs*. IMF_IP (P < 0.05) samples were observed; SM_NT was numerically higher than IMF_NT however statistical significance was not obtained (P = 0.08). In post-DUC 200K samples, significant differences were observed in particles/mL of SM_NT *vs*. IMF_NT (P < 0.01) and SM_AA vs IMF_AA (P < 0.05); statistical significance was not obtained between SM_IP *vs*. IMF_IP samples (Fig. 4b). This indicates that SM samples, both crude and pure, have higher EV like particles/mL of SM compared to IMF samples.

**Figure 4.**
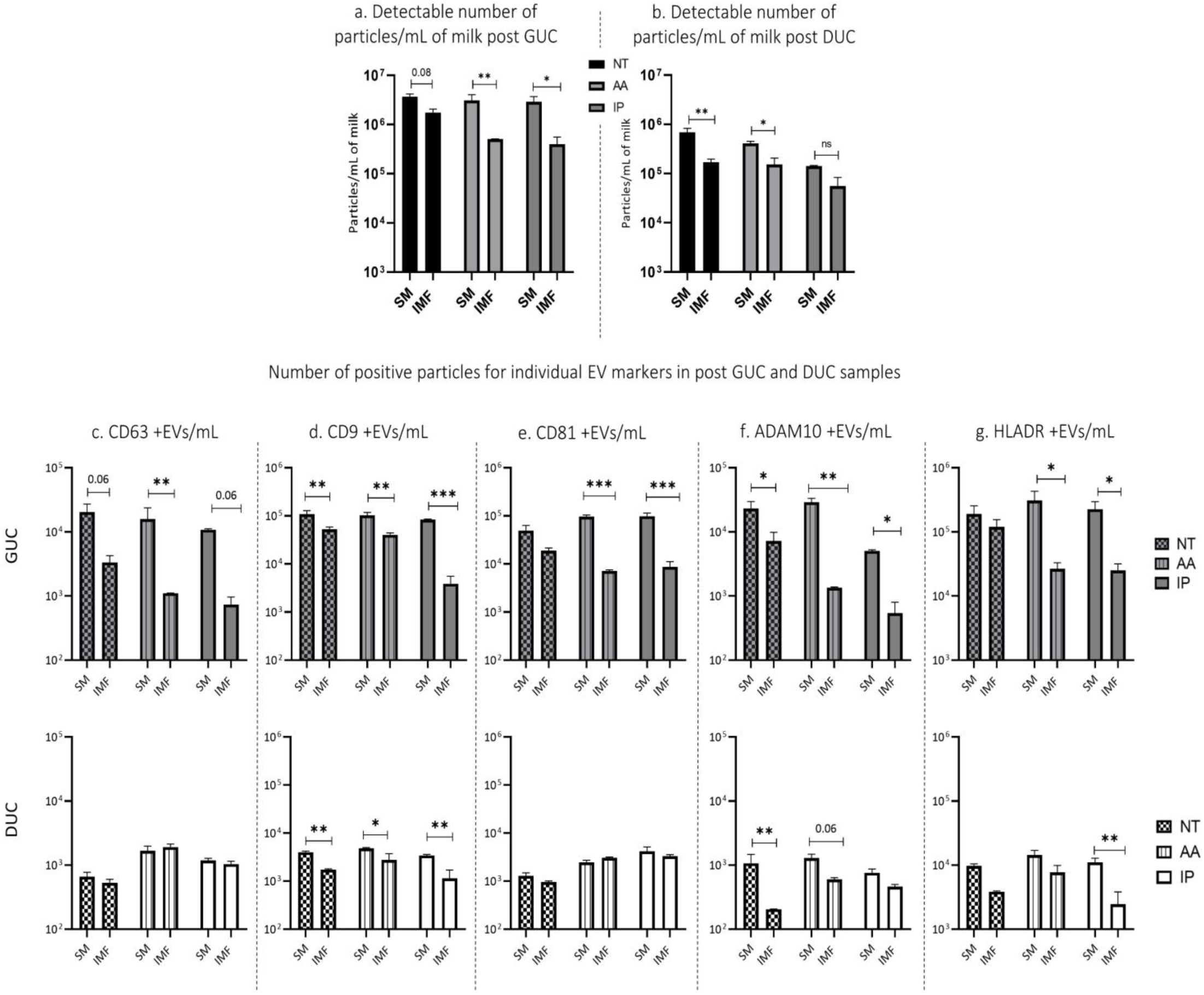
Imaging flow cytometry analysis of EV specific markers. Number of positive events (particles) in skim milk (SM) and infant milk formula (IMF) samples analysed by imaging flow cytometry (IFCM). All the samples were run at high magnification and low speed on Amnis ImageStream X Mk II imaging flow cytometer and data analysed on IDEAS software. The events were gated for low side scatter channel (SSC) signals and no signals on brightfield, based on EV size and events positive for these gating was acquired as objects (+EVs) in low SSC. The total number of detectable particles in gradient ultracentrifugation (GUC) samples are presented in a) and those from differential ultracentrifugation (DUC) samples are presented in c), data presented as mean and individual values ± SEM. The number of positive CD9, CD63, CD81, ADAM10 and HLADR particles in GUC samples are presented in b) and DUC samples are presented in d). The IDEAS software is used to further analyse the data, data presented as means and individual values ± SEM. A two-way Anova test was performed to compare differences in EVs in SM and IMF samples from each treatment groups; * indicates a significance value of P < 0.05. ** indicates a significance value of P < 0.01 and *** indicates a significance value of P < 0.001.

Fig. 4c, d, e, f & g present the number of +EVs/mL of sample for each individual EV specific marker. CD63: In post-GUC samples, all SM samples had higher number of CD63+ EVs/mL compared to IMF samples; the number of CD63+ EVs/mL in SM_AA were significantly higher compared to IMF_AA (P < 0.05), whereas numerically higher number of CD63 +EVs/mL with a trend towards statistical significance was observed between SM_NT and SM_IP compared to IMF_NT and IMF_IP (P = 0.06) samples, respectively. However, no significant differences (P > 0.05) between SM and IMF samples were observed post-DUC for the number of detected CD63+ EVs/mL.

CD9: In post-GUC samples, SM_NT, SM_AA and SM_IP samples had a significantly higher number of CD9+ EVs/mL compared to IMF_NT (P < 0.01), IMF_AA (P < 0.01) and IMF_IP (P < 0.001) samples, respectively. Similarly, for CD9+ EVs/mL in post-DUC samples, SM_NT. SM_AA and SM_IP samples had higher number of CD9+ EVs/mL compared to IMF_NT (P < 0.01), IMF_AA (P < 0.05) and IMF_IP (P < 0.01) samples, respectively.

CD81: In post-GUC samples, SM samples had significantly higher positive number of EVs/mL for CD81+ EVs/mL; SM_AA vs IMF_AA (P < 0.001) and SM_IP vs IMF_IP (P < 0.001). No significant differences between SM and IMF samples (P > 0.05) were observed in post-DUC samples.

ADAM10: Following the previous trend, the SM samples had a significantly higher number of ADAM10+ EVs/mL compared to the IMF samples; SM_NT vs IMF_NT (P < 0.05), SM_AA vs IMF_AA (P < 0.01) and SM_IP vs IMF_IP (P < 0.05). In post-DUC samples, significantly higher number of ADAM10+ EVs/mL were observed in SM_NT compared to IMF_NT (P < 0.01) samples.

HLADR: Lastly, differences in the HLADR+ EVs/mL in post-GUC SM samples were observed; SM_AA vs IMF_AA (P < 0.05) and SM_IP vs IMF_IP (P < 0.05). In post-DUC samples, SM_IP vs IMF_IP (P < 0.01) had higher number of HLADR+ EVs/mL; no significant difference was observed on HLADR+ EVs/mL in SM_NT vs IMF_NT and SM_AA vs IMF_AA samples.

## 4. Discussion

The aim of this study was to compare EVs present in SM and IMF. As IMF production involves harsh industrial processes, we hypothesised its EVs may be lost and/or compromised compared to those in SM. Comparisons were also made based on both milk pre-treatment and EV isolation methods to identify the optimal procedure to separate pure EVs from SM and IMF. Our results indicate that irrespective of the treatment, the concentration of EVs present in IMF are substantially compromised compared to SM and the lower amount of EVs or EV-like particles present in IMF do not carry EV specific markers. The results also suggest that both AA and IP pre-treatments remove milk proteins efficiently, and that the GUC technique is the optimum method for isolation of high yields of intact EVs without any effect on the EV morphology.

To perform detailed characterisation of milk derived EVs, pure EVs need to be separated. However, using complex biofluids in conjunction with ultracentrifugation typically leads to contamination of EVs with proteins (Langevin et al., 2019; Li et al., 2019). Interestingly, ultracentrifugation of milk leads to sedimentation of the dense casein micelles and all other proteins left in milk are called whey proteins (Fox, 2003). In our current study, we observed strong casein and whey protein bands by SDS-PAGE in samples that were collected after multiple ultracentrifugation steps of DUC in both SM and IMF samples; similar observations were also made for SM and IMF samples that underwent GUC. Previously, after milk fractionation by ultracentrifugation at 100,000 g, casein remnants were observed in milk serum because ultracentrifugation does not destroy the casein micelles and the smallest casein micelles are difficult to sediment (Jensen et al., 2012). This is why substantial consideration needs to be given towards removal of caseins due to their micelle forming property; casein micelles are approximately 150 nm in size and are naturally present in high concentrations in milk (Fox, 2001; O’Regan et al., 2009); both size and concentration overlapping with EVs.

In industry, casein is removed by isoelectric precipitation (Fox & Brodkorb, 2008) using acids that generate ‘acid-caseins’ (Atamer et al., 2017). Acidification of milk reduces its pH to the isoelectric point of casein (pH 4.6) resulting in irreversible destruction of the casein micelle structure and further coagulation and precipitation of the casein complex that can be sedimented at low speed centrifugation (Chinprahast et al., 2015; Fox & Brodkorb, 2008). In fact, a recent study investigated acidification effects on defatted raw bovine milk and observed some morphological changes due to acid treatment. However, the use of acid to remove milk proteins and separate large scale EVs remains their preferred method (Rahman et al., 2019). In our study, a combination of acid treatment and ultracentrifugation were applied to milk. To the best of our knowledge, this is a unique study where IMF samples are evaluated and compared to SM to identify how representative of SM is IMF, following the MISEV 2018 guidelines. The milk protein analysis on SDS-PAGE lead to exclusion of crude EV samples, based on the presence of casein protein bands, while pure EV samples were further characterised. Additionally, as a control, crude EV samples generated from NT group were included for all further analysis.

Further orthogonal characterisation was performed to analyse the EVs separated from SM and IMF and to explore the effects of pre-treatment on EV morphology or protein composition. Characterisation was performed by NTA, a technique widely used to estimate number of particles as well as the size of particles in a medium (Szatanek et al., 2017; Thery et al., 2018). Interestingly and as we hypothesised, post-GUC and -DUC, the number of particles/mL of IMF samples (all treatment groups) were significantly lower than in the corresponding SM samples, while no differences in the particle sizes were observed. Additionally, to rule out this being due to the effect of milk pre-treatment, we included the non-treated SM and IMF in our characterisation steps. Similar differences in non-treated SM and IMF samples were observed, where SM samples had higher concentration of EVs compared to IMF. This observation is in accordance to a very recent study where the authors compared IMF EVs to human and bovine milk and observed that while in IMF ∼10^10^ particles/mL were enumerated, these were mostly casein micelles or ‘exosome-sized-particles’ but not EVs (Leiferman et al., 2019).

While NTA is a widely used technique in EV research, it has certain limitations including its incapability to distinguish between particle sizes in a heterogeneous mixture being one (Witwer et al., 2013). Therefore, to further validate our observations, the EVs isolated from SM and IMF were imaged using TEM. The TEM analysis of GUC samples revealed that no negative effect of the pre-treatment or ultracentrifugation was observed on the EVs separated from SM. While the study by Rahman et al. (2019) observed certain morphological effects associated with milk acidification, here all the EVs derived from SM samples had a very smooth outer surface, round shaped and evenly distributed on the grid. This might be due to the differences in the methodology between the current study and that performed by Rahman et al. (2019). Here, compared to SM samples, however, GUC generated IMF samples were sparsely distributed across the grids and showed rough surfaces and shapes indicating to signs of disruption or disintegration. Interestingly in all the SM DUC pellets, very small particles (approximately 30 nm) were detected that did not have resemblance to the EVs that were observed in GUC samples. On contrary, no EV like structures were observed in IMF samples, with high amount of debris present in all IMF samples. The presence of high amount of debris in IMF might be related to the industrial processing during IMF production. Emami et al., (2018) reviewed the physicochemical degradation of protein formulations that undergo harsh spray drying, lyophilisation and heating steps; this possibly leads to high number of insoluble protein particles or aggregates that may pellet at the highest speed Additionally, these harsh industrial conditions also lead to disruption of EVs present in milk and hence no EV like structures were observed in IMF samples.

Further characterisation of the SM and IMF samples were done by immunoblotting for EV-positive markers. Two positive markers for EVs (tetraspanin CD63 and TSG101) and a negative milk EV marker (Actinin 4) were evaluated based on the MISEV 2018 guidelines (Table 3, (Thery et al., 2018)). In the GUC generated lysed samples, all SM samples, irrespective of their treatment, were positive for CD63 and TSG101 and negative for Actinin 4. Interestingly, IMF samples were associated with notably weaker signals indicating extremely low concentration of EVs present in the sample. Since the same amount of protein (35 µg) was loaded on the gel for each sample, lower density of bands indicates a very low EV to protein ratio indicating absence of EVs in the sample. A similar observation was also made in DUC samples where all IMF samples were again associated with faint bands for both CD63 and TSG101 compared to SM samples. These results, again, support our hypothesis that due to the use of harsh processing steps on milk in an industrial setting, EVs in the starting material are negatively affected. Of note, the doublet observed in some samples for TSG101 may be due to internal initiation at MET10 (VerPlank et al., 2001); whereas, in case of CD63, due to multiple glycosylation sites, bands within the range of 70 to 25 kDa are observed which is consistent with published literature (Metzelaar et al., 1991; Van Deun et al., 2014).

Lastly, the EVs separated from SM and IMF samples were analysed on AMNIS ImageStreamX Mark II Flow Cytometer following the recent published protocol (Ricklefs et al., 2019). Only events that had very low SSC signal and no brightfield signals were considered as a positive EV like particle and presented as particles/mL of milk. Following the observations from our previous characterisation steps, we observed that all SM samples, generated both by GUC and DUC, have a higher EV like particle counts compared to all IMF samples. Further, enumeration of surface markers on EVs that include tetraspanins CD9, CD81 and CD63, ADAM10 and HLADR were also carried out. In GUC generated samples, several significant differences were observed; all indicating that SM samples have higher EV specific surface markers compared to IMF samples, indicating again to the lower number of EVs present in IMF samples. In contrast, the SM samples generated by DUC recorded a higher CD9, ADAM10 and HLADR +EVs/mL compared to IMF samples. This observation may indicate that DUC in comparison to GUC does not separate pure EVs.

While this study is focussed primarily on comparing the differences between SM and IMF samples and we clearly demonstrated that IMF has lower amounts of EVs/EV-like particles, comparison between GUC vs DUC can also be made. The findings from this study are summarised in Table 2 and it is evident that GUC is a laborious process, with approximately 18 h required just for ultracentrifugation added to the time for sample preparation. However, the favourable aspect of this technique is the small volume of sample required compared to DUC. In fact, based on our NTA and IFCM results, GUC is associated with the highest yield of EVs compared to DUC; there is a log difference between the number of EVs detected in GUC compared to DUC. Additionally, the EV samples generated post-GUC, can be used in a multiple range of characterisation technique and generates highly reproducible data; apart from immunoblotting. In contrast, DUC yields high amount of visible pellet/suspension at the end of each centrifugation step, however, this suspension contains a high number of contaminant proteins that interferes with the further EV characterisation. The results from this study strongly indicate that a combination of AA/IP and GUC could be the best strategy to separate pure EVs from milk. In fact, GUC technique is a well-accepted method for isolation of EVs, especially the bottom-up approach, as previously reviewed (Witwer et al., 2013).

**Table 1.** Concentration of EVs separated from skim milk (SM) and infant milk formula (IMF) post gradient ultracentrifugation or differential ultracentrifugation, characterised using nanoparticle tracking analysis (NTA)

| **a) Gradient Ultracentrifugation** |  |  |  |  |  |
| --- | --- | --- | --- | --- | --- |
| **Concentration**<br>**(Particles/mL of starting material)** | **Skim milk** |  | **Infant milk formula** |  | **P value** |
| **Non-treated (NT)** | $4.35 \times 10^{10} \pm 1.40 \times 10^{10}$ | | $6.13 \times 10^{09} \pm 8.17 \times 10^{08}$ | | 0.05 |
| **Acetic acid treated (AA)** | $1.39 \times 10^{10} \pm 8.25 \times 10^{09}$ | | $2.82 \times 10^{09} \pm 2.33 \times 10^{09}$ | | ns |
| **Isoelectric precipitation (IP)** | $2.16 \times 10^{10} \pm 2.47 \times 10^{09}$ | | $3.22 \times 10^{09} \pm 1.45 \times 10^{09}$ | | 0.01 |
| **Size (in nm)** | Mean | Mode | Mean | Mode |  |
| **Non-treated (NT)** | 147.00 ± 8.39 | 113.60 ± 10.34 | 166.15 ± 2.75 | 140.45 ± 1.25 | ns |
| **Acetic acid treated (AA)** | 159.83 ± 5.55 | 135.46 ± 5.28 | 180.16 ± 16.77 | 133.53 ± 16.22 | ns |
| **Isoelectric precipitation (IP)** | 162.90 ± 13.51 | 141.03 ± 11.93 | 152.40 ± 45.10 | 109.50 ± 18.80 | ns |
| **b) Differential Ultracentrifugation** |  |  |  |  |  |
| **Concentration**<br>**(Particles/mL of starting material)** | **Skim milk** |  | **Infant milk formula** |  | **P value** |
| **Non-treated (NT)** | $1.97 \times 10^{10} \pm 8.90 \times 10^{09}$ | | $4.65 \times 10^{07} \pm 2.50 \times 10^{07}$ | | 0.04 |
| **Acetic acid treated (AA)** | $2.78 \times 10^{09} \pm 1.00 \times 10^{09}$ | | $1.69 \times 10^{08} \pm 3.30 \times 10^{07}$ | | 0.03 |
| **Isoelectric precipitation (IP)** | $3.65 \times 10^{09} \pm 5.43 \times 10^{08}$ | | $4.66 \times 10^{08} \pm 2.02 \times 10^{08}$ | | 0.01 |
| **Size (in nm)** | Mean | Mode | Mean | Mode |  |
| **Non-treated (NT)** | 173.60 ± 29.20 | 123.40 ± 60.00 | 180.40 ± 7.50 | 152.50 ± 50.10 | ns |
| **Acetic acid treated (AA)** | 150.10 ± 16.40 | 146.90 ± 47.33 | 149.28 ± 18.37 | 137.30 ± 68.22 | ns |
| **Isoelectric precipitation (IP)** | 149.23 ± 23.50 | 130.81 ± 46.11 | 184.19 ± 15.63 | 117.10 ± 67.70 | ns |

ns = Not significant

**Table 2.** Summary of the techniques used for separation of milk EVs and the characterisation of EVs.

| Milk type | Pre-treatment | EV separation method | Vol. of sample used (mL) | Time to separate EV (h) | Immuno-blotting | EV size (nm) | EV concentration /mL of milk | TEM | IFCM |
| --- | --- | --- | --- | --- | --- | --- | --- | --- | --- |
| SM | NT | GUC | 2.33 | > 18 | (?) | 146.4 ± 20.36 | 10<sup>12</sup> | + | 10<sup>6</sup> |
|  | AA | GUC | 2.33 | > 18 | (?) | 166.7 ± 4.80 | 10<sup>12</sup> | + | 10<sup>6</sup> |
|  | IP | GUC | 2.33 | > 18 | (?) | 155.3 ± 8.48 | 10<sup>11</sup> | + | 10<sup>6</sup> |
| IMF | NT | GUC | 2.33 | > 18 | (?) | 176.3 ± 39.88 | 10<sup>11</sup> | + | 10<sup>6</sup> |
|  | AA | GUC | 2.33 | > 18 | (?) | 169.1 ± 14.84 | 10<sup>11</sup> | + | 10<sup>5</sup> |
|  | IP | GUC | 2.33 | > 18 | (?) | 134.3 ± 39.45 | 10<sup>10</sup> | + | 10<sup>5</sup> |
| SM | NT | DUC | 38 | < 6.25 | (?) | 150.8 ± 63.00 | 10<sup>11</sup> | + (high debris) | 10<sup>5</sup> |
|  | AA | DUC | 38 | < 6.25 | (?) | 168.9 ± 50.10 | 10<sup>10</sup> | + (high debris) | 10<sup>5</sup> |
|  | IP | DUC | 38 | < 6.25 | (?) | 157.6 ± 47.30 | 10<sup>11</sup> | + | 10<sup>5</sup> |
| IMF | NT | DUC | 38 | < 6.25 | (?) | 188.9 ± 68.20 | 10<sup>9</sup> | + (high debris) | 10<sup>5</sup> |
|  | AA | DUC | 38 | < 6.25 | (?) | 160.0 ± 46.00 | 10<sup>9</sup> | + (high debris) | 10<sup>5</sup> |
|  | IP | DUC | 38 | < 6.25 | (?) | 178.3 ± 67.70 | 10<sup>10</sup> | + (high debris) | 10<sup>4</sup> |

SM skim milk, IMF infant milk formula
+ vesicle like structures detected
(?) samples could be analysed without technical limitations

Therefore, returning to our question of how representative IMF is of SM, we can clearly state that IMF contains significantly fewer EVs/EV-like particles which have compromised morphologies and do not carry EV specific markers. Additionally, based on the current results we can also conclude that methodology that includes IP treatment and GUC although takes longer uses comparatively less amount of starting material and the separated EVs can be used for techniques such as NTA, TEM and IFCM.

## Acknowledgements

This work was supported by the Department of Agriculture, Food & Marine, Ireland under Grant [17/F/234]. IFCM was performed on instrumentation in the Flow Cytometry Facility, Trinity Biomedical Sciences Institute, funded by Science Foundation Ireland.

## References

Admyre, C., Johansson, S. M., Qazi, K. R., Filen, J. J., Lahesmaa, R., Norman, M., Neve, E. P., Scheynius, A., & Gabrielsson, S. (2007). Exosomes with immune modulatory features are present in human breast milk. J Immunol, 179(3), 1969–1978.

Arntz, O. J., Pieters, B. C., Oliveira, M. C., Broeren, M. G., Bennink, M. B., de Vries, M., van Lent, P. L., Koenders, M. I., van den Berg, W. B., van der Kraan, P. M., et al. (2015). Oral administration of bovine milk derived extracellular vesicles attenuates arthritis in two mouse models. Molecular Nutrition and Food Research, 59(9), 1701–1712. doi: 10.1002/mnfr.201500222

Atamer, Z., Post, A. E., Schubert, T., Holder, A., Boom, R. M., & Hinrichs, J. J. I. d. j. (2017). Bovine β-casein: Isolation, properties and functionality. A review. International dairy journal, 66, 115–125.

Benmoussa, A., Lee, C. H. C., Laffont, B., Savard, P., Laugier, J., Boilard, E., Gilbert, C., Fliss, I., & Provost, P. (2016). Commercial Dairy Cow Milk microRNAs Resist Digestion under Simulated Gastrointestinal Tract Conditions. The Journal of nutrition, 146(11), 2206–2215.

Benmoussa, A., Ly, S., Shan, S. T., Laugier, J., Boilard, E., Gilbert, C., & Provost, P. (2017). A subset of extracellular vesicles carries the bulk of microRNAs in commercial dairy cow’s milk. Journal of Extracellular Vesicles, 6(1), 1401897.

Benmoussa, A., & Provost, P. (2019). Milk microRNAs in health and disease. Comprehensive Reviews in food science and food safety, 18(3), 703–722.

Burke, N., Zacharski, K. A., Southern, M., Hogan, P., Ryan, M. P., & Adley, C. C. (2018). The Dairy Industry: Process, Monitoring, Standards, and Quality. In Descriptive Food Science: IntechOpen.

Chinprahast, N., Subhimaros, S., & Pattorn, S. J. I. F. R. J. (2015). Heat-acid coagulation of market-returned UHT milk using various coagulants and calcium chloride. International Food Research Journal, 22(3).

Colombo, M., Raposo, G., & Théry, C. (2014). Biogenesis, secretion, and intercellular interactions of exosomes and other extracellular vesicles. Annual review of cell and developmental biology, 30, 255–289.

Emami, F., Vatanara, A., Park, E. J., & Na, D. H. (2018). Drying technologies for the stability and bioavailability of biopharmaceuticals. Pharmaceutics, 10(3), 131.

Fox, P. (2001). Milk proteins as food ingredients. International Journal of Dairy Technology, 54(2), 41–55.

Fox, P. (2003). Milk proteins: general and historical aspects. In Advanced Dairy Chemistry—1 Proteins (pp. 1–48): Springer.

Fox, P., & Brodkorb, A. (2008). The casein micelle: Historical aspects, current concepts and significance. International dairy journal, 18(7), 677–684.

Gardiner, C., Vizio, D. D., Sahoo, S., Théry, C., Witwer, K. W., Wauben, M., & Hill, A. F. (2016). Techniques used for the isolation and characterization of extracellular vesicles: results of a worldwide survey. Journal of Extracellular Vesicles, 5(1), 32945.

Gathercole, J., Reis, M. G., Agnew, M., Reis, M. M., Humphrey, R., Harris, P., Clerens, S., Haigh, B., & Dyer, J. M. (2017). Molecular modification associated with the heat treatment of bovine milk. International dairy journal, 73, 74–83.

Gonzales, P. A., Zhou, H., Pisitkun, T., Wang, N. S., Star, R. A., Knepper, M. A., & Yuen, P. S. (2010). Isolation and purification of exosomes in urine. In The Urinary Proteome (pp. 89–99): Springer.

Hata, T., Murakami, K., Nakatani, H., Yamamoto, Y., Matsuda, T., & Aoki, N. (2010). Isolation of bovine milk-derived microvesicles carrying mRNAs and microRNAs. Biochem Biophys Res Commun, 396(2), 528–533. doi: 10.1016/j.bbrc.2010.04.135

Izumi, H., Kosaka, N., Shimizu, T., Sekine, K., Ochiya, T., & Takase, M. (2012). Bovine milk contains microRNA and messenger RNA that are stable under degradative conditions. J Dairy Sci, 95(9), 4831–4841. doi: 10.3168/jds.2012-5489

Jensen, H. B., Poulsen, N. A., Møller, H. S., Stensballe, A., & Larsen, L. B. (2012). Comparative proteomic analysis of casein and whey as prepared by chymosin-induced separation, isoelectric precipitation or ultracentrifugation. Journal of Dairy Research, 79(4), 451–458.

Kowal, J., Arras, G., Colombo, M., Jouve, M., Morath, J. P., Primdal-Bengtson, B., Dingli, F., Loew, D., Tkach, M., & Théry, C. (2016). Proteomic comparison defines novel markers to characterize heterogeneous populations of extracellular vesicle subtypes. Proceedings of the National Academy of Sciences, 113(8), E968–E977.

Kusuma, R. J., Manca, S., Friemel, T., Sukreet, S., Nguyen, C., & Zempleni, J. (2016). Human vascular endothelial cells transport foreign exosomes from cow’s milk by endocytosis. American Journal of Physiology-Cell Physiology, 310(10), C800–807. doi: 10.1152/ajpcell.00169.2015

Langevin, S. M., Kuhnell, D., Orr-Asman, M. A., Biesiada, J., Zhang, X., Medvedovic, M., & Thomas, H. E. (2019). Balancing yield, purity and practicality: a modified differential ultracentrifugation protocol for efficient isolation of small extracellular vesicles from human serum. RNA Biology, 16(1), 5–12. doi: 10.1080/15476286.2018.1564465

Lasser, C., Alikhani, V. S., Ekstrom, K., Eldh, M., Paredes, P. T., Bossios, A., Sjostrand, M., Gabrielsson, S., Lotvall, J., & Valadi, H. (2011). Human saliva, plasma and breast milk exosomes contain RNA: uptake by macrophages. J Transl Med, 9, 9. doi: 10.1186/1479-5876-9-9

Leiferman, A., Shu, J., Upadhyaya, B., Cui, J., & Zempleni, J. (2019). Storage of extracellular vesicles in human milk, and microRNA profiles in human milk exosomes and infant formulas. Journal of pediatric gastroenterology and nutrition, 69(2), 235–238.

Li, J., He, X., Deng, Y., & Yang, C. (2019). An Update on Isolation Methods for Proteomic Studies of Extracellular Vesicles in Biofluids. Molecules, 24(19). doi: 10.3390/molecules24193516

Manca, S., Upadhyaya, B., Mutai, E., Desaulniers, A. T., Cederberg, R. A., White, B. R., & Zempleni, J. (2018). Milk exosomes are bioavailable and distinct microRNA cargos have unique tissue distribution patterns. Sci Rep, 8(1), 11321. doi: 10.1038/s41598-018-29780-1

Metzelaar, M., Wijngaard, P., Peters, P., Sixma, J. J., Nieuwenhuis, H. K., & Clevers, H. (1991). CD63 antigen. A novel lysosomal membrane glycoprotein, cloned by a screening procedure for intracellular antigens in eukaryotic cells. Journal of Biological Chemistry, 266(5), 3239–3245.

Munagala, R., Aqil, F., Jeyabalan, J., & Gupta, R. C. (2016). Bovine milk-derived exosomes for drug delivery. Cancer Letter, 371(1), 48–61. doi: 10.1016/j.canlet.2015.10.020

Musante, L., Saraswat, M., Ravidà, A., Byrne, B., & Holthofer, H. (2013). Recovery of urinary nanovesicles from ultracentrifugation supernatants. Nephrology Dialysis Transplantation, 28(6), 1425–1433.

O’Brien, K., Rani, S., Corcoran, C., Wallace, R., Hughes, L., Friel, A. M., McDonnell, S., Crown, J., Radomski, M. W., & O’Driscoll, L. (2013). Exosomes from triple-negative breast cancer cells can transfer phenotypic traits representing their cells of origin to secondary cells. European Journal of Cancer, 49(8), 1845–1859. doi: 10.1016/j.ejca.2013.01.017

O’Regan, J., Ennis, M., & Mulvihill, D. (2009). Milk proteins. In Handbook of hydrocolloids (pp. 298–358): Elsevier.

Rahman, M. M., Shimizu, K., Yamauchi, M., Takase, H., Ugawa, S., Okada, A., & Inoshima, Y. (2019). Acidification effects on isolation of extracellular vesicles from bovine milk. PLoS One, 14(9).

Reinhardt, T. A., Lippolis, J. D., Nonnecke, B. J., & Sacco, R. E. (2012). Bovine milk exosome proteome. Journal of proteomics, 75(5), 1486–1492. doi: 10.1016/j.jprot.2011.11.017

Reinhardt, T. A., Sacco, R. E., Nonnecke, B. J., & Lippolis, J. D. (2013). Bovine milk proteome: quantitative changes in normal milk exosomes, milk fat globule membranes and whey proteomes resulting from Staphylococcus aureus mastitis. Journal of proteomics, 82, 141–154. doi: 10.1016/j.jprot.2013.02.013

Ricklefs, F. L., Maire, C. L., Reimer, R., Dührsen, L., Kolbe, K., Holz, M., Schneider, E., Rissiek, A., Babayan, A., Hille, C., et al. (2019). Imaging flow cytometry facilitates multiparametric characterization of extracellular vesicles in malignant brain tumours. Journal of Extracellular Vesicles, 8(1), 1588555. doi: 10.1080/20013078.2019.1588555

Samuel, M., Chisanga, D., Liem, M., Keerthikumar, S., Anand, S., Ang, C.-S., Adda, C. G., Versteegen, E., Jois, M., & Mathivanan, S. (2017). Bovine milk-derived exosomes from colostrum are enriched with proteins implicated in immune response and growth. Sci Rep, 7(1), 1–10.

Somiya, M., Yoshioka, Y., & Ochiya, T. (2018). Biocompatibility of highly purified bovine milk-derived extracellular vesicles. Journal of Extracellular Vesicles, 7(1), 1440132. doi: 10.1080/20013078.2018.1440132

Sun, J., Aswath, K., Schroeder, S. G., Lippolis, J. D., Reinhardt, T. A., & Sonstegard, T. S. (2015). MicroRNA expression profiles of bovine milk exosomes in response to Staphylococcus aureus infection. BMC Genomics, 16, 806. doi: 10.1186/s12864-015-2044-9

Sun, Q., Chen, X., Yu, J., Zen, K., Zhang, C. Y., & Li, L. (2013). Immune modulatory function of abundant immune-related microRNAs in microvesicles from bovine colostrum. Protein Cell, 4(3), 197–210. doi: 10.1007/s13238-013-2119-9

Szatanek, R., Baj-Krzyworzeka, M., Zimoch, J., Lekka, M., Siedlar, M., & Baran, J. (2017). The Methods of Choice for Extracellular Vesicles (EVs) Characterization. International Journal of Molecular Sciences, 18(6). doi: 10.3390/ijms18061153

Thery, C., Witwer, K. W., Aikawa, E., Alcaraz, M. J., Anderson, J. D., Andriantsitohaina, R., Antoniou, A., Arab, T., Archer, F., Atkin-Smith, G. K., et al. (2018). Minimal information for studies of extracellular vesicles 2018 (MISEV2018): a position statement of the International Society for Extracellular Vesicles and update of the MISEV2014 guidelines. Journal of Extracellular Vesicles, 7(1), 1535750. doi: 10.1080/20013078.2018.1535750

Van Deun, J., Mestdagh, P., Agostinis, P., Akay, Ö., Anand, S., Anckaert, J., Martinez, Z. A., Baetens, T., Beghein, E., & Bertier, L. (2017). EV-TRACK: transparent reporting and centralizing knowledge in extracellular vesicle research. Nature methods, 14(3), 228.

Van Deun, J., Mestdagh, P., Sormunen, R., Cocquyt, V., Vermaelen, K., Vandesompele, J., Bracke, M., De Wever, O., & Hendrix, A. (2014). The impact of disparate isolation methods for extracellular vesicles on downstream RNA profiling. Journal of Extracellular Vesicles, 3(1), 24858.

van Herwijnen, M. J., Zonneveld, M. I., Goerdayal, S., Nolte-’t Hoen, E. N., Garssen, J., Stahl, B., Maarten Altelaar, A. F., Redegeld, F. A., & Wauben, M. H. (2016). Comprehensive Proteomic Analysis of Human Milk-derived Extracellular Vesicles Unveils a Novel Functional Proteome Distinct from Other Milk Components. Molecular & Cellular Proteomics 15(11), 3412–3423. doi: 10.1074/mcp.M116.060426

van Herwijnen, M. J. C., Driedonks, T. A. P., Snoek, B. L., Kroon, A. M. T., Kleinjan, M., Jorritsma, R., Pieterse, C. M. J., Hoen, E., & Wauben, M. H. M. (2018). Abundantly Present miRNAs in Milk-Derived Extracellular Vesicles Are Conserved Between Mammals. Frontiers in Nutrition, 5, 81. doi: 10.3389/fnut.2018.00081

Vaswani, K., Koh, Y. Q., Almughlliq, F. B., Peiris, H. N., & Mitchell, M. D. (2017). A method for the isolation and enrichment of purified bovine milk exosomes. Reprod Biol, 17(4), 341–348. doi: 10.1016/j.repbio.2017.09.007

VerPlank, L., Bouamr, F., LaGrassa, T. J., Agresta, B., Kikonyogo, A., Leis, J., & Carter, C. A. (2001). Tsg101, a homologue of ubiquitin-conjugating (E2) enzymes, binds the L domain in HIV type 1 Pr55Gag. Proceedings of the National Academy of Sciences, 98(14), 7724–7729.

Visnovitz, T., Osteikoetxea, X., Sodar, B. W., Mihaly, J., Lorincz, P., Vukman, K. V., Toth, E. A., Koncz, A., Szekacs, I., Horvath, R., et al. (2019). An improved 96 well plate format lipid quantification assay for standardisation of experiments with extracellular vesicles. Journal of Extracellular Vesicles, 8(1), 1565263. doi: 10.1080/20013078.2019.1565263

Witwer, K. W., Buzas, E. I., Bemis, L. T., Bora, A., Lasser, C., Lotvall, J., Nolte-’t Hoen, E. N., Piper, M. G., Sivaraman, S., Skog, J., et al. (2013). Standardization of sample collection, isolation and analysis methods in extracellular vesicle research. Journal of Extracellular Vesicles, 2. doi: 10.3402/jev.v2i0.20360

Wolf, T., Baier, S. R., & Zempleni, J. (2015). The Intestinal Transport of Bovine Milk Exosomes Is Mediated by Endocytosis in Human Colon Carcinoma Caco-2 Cells and Rat Small Intestinal IEC-6 Cells. Journal of Nutrition, 145(10), 2201–2206. doi: 10.3945/jn.115.218586

Yamada, T., Inoshima, Y., Matsuda, T., & Ishiguro, N. (2012). Comparison of methods for isolating exosomes from bovine milk. Journal of Veterinary Medical Science, 74(11), 1523–1525.

Yamauchi, M., Shimizu, K., Rahman, M., Ishikawa, H., Takase, H., Ugawa, S., Okada, A., & Inoshima, Y. (2018a). Efficient method for isolation of exosomes from raw bovine milk. Drug Dev Ind Pharm, 1–6. doi: 10.1080/03639045.2018.1539743

Yamauchi, M., Shimizu, K., Rahman, M., Ishikawa, H., Takase, H., Ugawa, S., Okada, A., & Inoshima, Y. (2018b). Efficient method for isolation of exosomes from raw bovine milk. Drug development and industrial pharmacy, 45(3), 359–364. doi: 10.1080/03639045.2018.1539743

Zonneveld, M. I., Brisson, A. R., van Herwijnen, M. J., Tan, S., van de Lest, C. H., Redegeld, F. A., Garssen, J., Wauben, M. H., & Nolte-’t Hoen, E. N. (2014). Recovery of extracellular vesicles from human breast milk is influenced by sample collection and vesicle isolation procedures. Journal of Extracellular Vesicles, 3(1), 24215.

